# Reduced Folate Carrier 1 (RFC1/Slc19a1) Suppression Exacerbates Blood-Brain Barrier Breakdown in Experimental Ischemic Stroke in Adult Mice

**DOI:** 10.1101/2024.10.28.620539

**Authors:** Gokce Gurler, Dilan Bozanoglu, Christelle Leon, Nevin Belder, Melike Sever Bahcekapili, Radu Bolbos, Hulya Karatas, Marlene Wiart, Fabien Chauveau, Muge Yemisci

**Affiliations:** Institute of Neurological Sciences and Psychiatry, Hacettepe University, Ankara, Turkey; Univ. Lyon, CarMeN laboratory, IRIS team, INSERM, INRAE, Bron, France; Biotechnology Institute, Ankara University, Ankara, Turkey; CERMEP-Imagerie du Vivant, Bron, France; Neuroscience and Neurotechnology Center of Excellence (NÖROM) Ankara, Turkey; CNRS, 69100 Villeurbanne, France; Univ. Lyon, Lyon Neuroscience Research Center, BIORAN team, CNRS, INSERM, Bron, France; Faculty of Medicine, Department of Neurology, Hacettepe University, Ankara, Turkey; Graduate School of Health Sciences, Hacettepe University, Ankara, Turkey

**Author notes:** Corresponding Author: Prof. Muge Yemisci (MD, PhD) Institute of Neurological Sciences and Psychiatry, Hacettepe University, Ankara, Turkey Faculty of Medicine, Department of Neurology, Hacettepe University, Ankara, Turkey Neuroscience and Neurotechnology Center of Excellence (NÖROM) Ankara, Turkey. These authors contributed equally to this work (Fabien Chauveau, Muge Yemisci).

**Keywords:** blood-brain barrier, ischemic stroke, magnetic resonance imaging, RFC1, SLC19A1, Slc19a1

## Abstract

The Reduced Folate Carrier 1 (RFC1), also called solute carrier family 19 member 1 (SLC19A1/SLC19a1), is recognized for transporting folates across the blood-brain barrier (BBB). RFC1 has recently been defined as a hypoxia-immune related gene whose expression levels were induced by acute retinal ischemia, suggesting that RFC1 may have a role in the response of the brain to ischemic injury. Despite a recent human meta-analysis suggesting an association between certain RFC1 polymorphisms and the risk of silent brain infarctions, preclinical evidence concerning the potential role of RFC1 in acute ischemic stroke has yet to be presented. To investigate this, we first characterized RFC1 protein expression in mouse microvessels and pericytes which play significant roles in stroke pathophysiology. Then, we examined the temporal (1-h, 24-h, and 48-h) and spatial (infarct, periinfarct, contralateral) expression of RFC1 protein in the intraluminal transient middle cerebral artery occlusion mouse model. Finally, we knocked down RFC1 protein with RFC1-siRNA in the potential periinfarct region before induction of ischemia and investigated BBB integrity and infarct size *in vivo* via 7T-MRI. Moreover, we utilized a pharmacological modulation-methotrexate, a non-covalent inhibitor of RFC1- to further investigate the role of RFC1 in maintaining BBB integrity. Our study revealed that, i) RFC1 protein levels were dynamic throughout the acute phases of ischemic stroke, ii) RFC1 suppression aggravated the BBB leakage during ischemia. These results emphases the role of RFC1 in the pathophysiology of ischemic stroke and supports the evidence from human studies.

## BACKGROUND

Cerebrovascular diseases represent a global health challenge due their severe impact and often poor prognosis. Acute ischemic stroke, caused by the abrupt blockage of the arteries, is the leading cause of death and disability among cerebrovascular diseases. One of the major factors determining functional outcome after acute ischemic stroke is the blood-brain barrier (BBB) disruption aggravating neuronal injury besides the reduced blood flow (1,2). Despite decades of research, dissecting the elements that play important role in BBB disruption in ischemic stroke pathophysiology is still a work in progress.

The BBB is an essential barrier within the brain’s microvasculature, protecting the parenchyma from harmful substances and regulating the optimal concentration of essential nutrients under physiological conditions. The Reduced Folate Carrier 1 (RFC1), also known as solute carrier family 19 member 1 (SLC19A1/SLC19a1), is harbored in the brain microvessels and is a major transporter of reduced folates. Folates, belonging to the B9 group of vitamins, play crucial roles in maintaining optimal Central Nervous System (CNS) function such as DNA methylation, replication, repair, amino acid metabolism, neurotransmitter modulation, and phospholipid synthesis (3,4). Beyond its role as a folate carrier in the BBB, our recent studies demonstrated that RFC1 protein is significant in maintaining the integrity of the inner blood-retina barrier (BRB), which is the extension of the BBB anatomically, functionally, and in response to insults, in healthy retinas (5). Furthermore, we revealed the role of RFC1 in acute retinal ischemia, as evidenced by increased retinal RFC1 protein levels following 1-h acute retinal ischemia in mice.

The polymorphisms of RFC1 have also been linked to cerebrovascular diseases such as ischemic stroke and silent brain infarction in a human study (6) Moreover, RFC1 is recognized as a hypoxia-sensitive gene, indicating its vulnerability to oxidative stress modifications (7). These findings combined with our findings in retina underscore the potential significance of RFC1 in the pathogenesis of stroke and related conditions. Besides, our recent studies, involving mice brain sections, whole-mount retinas, and retinal and cerebral microvessel isolates, demonstrated RFC1 protein expression in retinal and brain pericytes which are crucial cells within the neurovascular unit (8,9). Pericytes are involved in numerous functions including the maintenance of vascular stability, modulation of neuroinflammation, participation in angiogenesis and neurovascular coupling as well as playing a major role in the pathophysiology of acute ischemic stroke. (10). While our findings in ischemic retinas were a first step linking acute ischemia and RFC1, this hypothesis lacks direct preclinical evidence in the pathophysiology of ischemic stroke of the brain. Furthermore, the role of RFC1 in acute cerebral ischemia remains unexplored, as there are no previous investigations into its involvement despite clinical evidence.

The present study aims to investigate the potential involvement of RFC1 in the pathophysiology of acute cerebral ischemia. We utilized a custom-designed RFC1-targeted short interfering RNA (siRNA) to silence the RFC1 gene *in vivo*. We studied the effect of RFC1 silencing on cerebral ischemia/recanalization by using transient middle cerebral artery occlusion (MCAo) model in mice. The functional consequences of this modulation on BBB integrity were explored through longitudinal *in vivo* 7T magnetic resonance imaging (MRI). Besides, to modify RFC1 levels pharmacologically we used methotrexate (MTX), a drug which is used clinically as an antineoplastic and antirheumatic agent, and functions as a non-covalent (competitive) inhibitor of RFC1 (11). The reduction of RFC1 levels enhanced BBB permeability. Thus, in this study we share our findings underscoring the immediate role of RFC1 in maintaining BBB integrity in both health and acute cerebral ischemic conditions.

## MATERIALS AND METHODS

### Animals

Adult male and female Swiss albino mice (25-35 g) were used for the experiments. All the procedures were approved by Hacettepe University Animal Experimentations Local Ethics Board (No: 2019/13-06 and 2021/03-19), and the MRI study was approved by the local ethical review board (“CELYNE”, registration code: C2EA-42), authorized by the French Ministry of Higher Education and Research (APAFIS #36695-2022051017481524 v1), and was in accordance with European directives on the protection and use of laboratory animals (Council Directive 2010/63/UE, French decree 2013-118). Animal housing, care, and the experimental procedures were done in accordance with institutional guidelines and reporting complies with ARRIVE guidelines (Animal Research: Reporting of In Vivo Experiments) (12). Same sex littermates of male and female mice were kept together in conventional cages (Type II open polycarbonate cage) in rooms with a 12-h light–dark cycle, controlled temperature (22±2 °C) and humidity (50-60%). All mice were provided with ad libitum standard mouse diet and drinking water. The experimental groups matched control groups, and the numbers of mice per groups are given in Table 1. Animals were allocated to groups randomly and without regarding to the order of treatments and measurements. The allocation or conduct of the experiments or data analysis were performed by different experimenters who were blinded to the groups. For the surgical procedures mentioned below, mice were anesthetized with isoflurane (4-5% induction dose, followed by 1.5-2% maintenance). Corneal reflex and hind-paw nociceptive reflex sensitivity were assessed periodically for depth of anesthesia. Air/O_2_ flow rate was maintained around 2 L/min via a nose cone. Body temperature was monitored with a rectal probe and maintained at 37.0±0.2 °C by a homeothermic blanket control unit (Harvard Apparatus, USA). Pulse rate and oxygen saturation were monitored with an oximeter (The LifeSense® VET Pulse Oximeter, Nonin Medical Inc., USA) from the right lower limb. Analgesia was administered subcutaneously (0.05 mg/kg buprenorphine) prior to surgery, and the surgical incision was locally anesthetized with lidocaine.

**Table 1.**
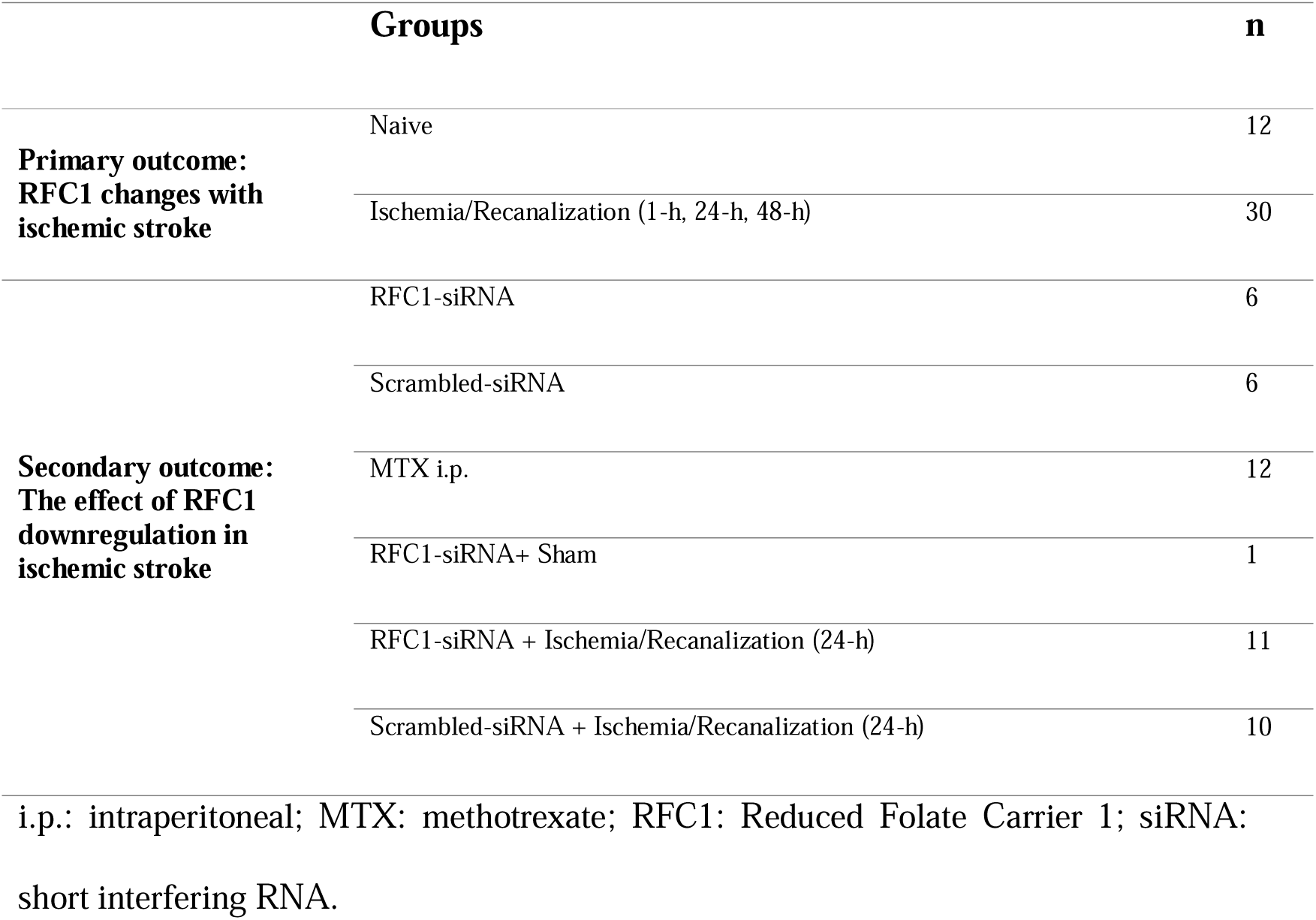
Table of groups and number of mice used for the experiments.

### Transient Focal Cerebral Ischemia

Intraluminal monofilament technique was used for right proximal middle cerebral artery occlusion (MCAo) and recanalization as previously described (13,14). Briefly, deeply anesthetized mouse was placed under a surgical microscope in supine position. The right common, external, and internal carotid arteries were dissected through a midline neck incision. The common and external carotid arteries were ligated with 6-0 nylon suture proximally. Through a small incision 1 mm proximal to carotid bifurcation on carotid artery, silicon rubber-coated monofilament (Doccol, 602412PK10) was introduced into the internal carotid artery.

For the animals which underwent MRI, the caudal tail was catheterized to inject contrast agent, and the success of occlusion was confirmed by MR angiography performed during the occlusion (see below).

For the other animals, the success of occlusion and recanalization was confirmed by measuring the regional cerebral blood flow with laser-Doppler flowmetry (PeriFlux 6000, PERIMED, Sweden) via a flexible probe (PF-318 of PeriFlux PF 2B, Perimed Jarfalla, Sweden) placed over the skull (2 mm posterior, 6 mm lateral to the bregma). After obtaining a stable preischemic relative cerebral blood flow (rCBF) for 10 min, the MCA was occluded by advancing the filament 10 mm distal to the carotid bifurcation. Observing ∼90% drop in rCBF in laser-Doppler flowmetry confirmed the total occlusion of MCA. rCBF was continuously monitored during ischemia (90 min) and the first 10 min of recanalization. After 90 min, the filament was drawn for recanalization for 1 h, 24 h and 48 h. Sham surgery was performed with all surgical steps except the introduction of the filament. Mice were placed on a heating blanket until they will fully recover from anesthesia. The animals that showed hemodynamic instability, subarachnoid hemorrhage, incomplete occlusion or died during surgery were excluded. Animals who had complete occlusion of the MCA validated by laser doppler flowmetry or TOF-MR angiography were included in experimental groups. Neurological motor examination was performed 24-h after the MCAo according to the Modified Bederson score as grade 0: no observable neurologic deficit (normal), grade 1: forelimb flexion, grade 2: forelimb flexion and decreased resistance to lateral push, grade 3: circling, grade 4: circling and spinning around the cranial-caudal axis, grade 5: no spontaneous movement contralateral side at rest or no spontaneous motor activity (severe) (15).

### Immunohistochemical Studies

Brains, extracted from transcardially perfused mice with 4% paraformaldehyde (PFA) solution, were immersed in 4% PFA for 24-h followed by 30% sucrose (in PBS) for 2 days for cryopreservation. Brain sections of 20 µm thick were used. Also, brain microvessels were isolated (see below) and stained separately. The sections or isolated microvessels were incubated in Tris-buffered saline (TBS) containing 0.3% Triton-X for 30 min for permeabilization followed by blocking with 10% normal goat serum (NGS) for 1 h at room temperature. Next, they were incubated overnight at + 4°C with primary antibodies against RFC1 (SLC19A1, MyBioSource MBS9134642 and Sigma-Aldrich AV44167; both produced in rabbit); for mural cells against PDGFR-β (R&D Systems, AF1042), CD13 (Acris Antibodies, AM26636AF-N); for endothelium CD31 (BD Bioscience, 550274). Cy3-conjugated anti-mouse IgG (Jackson ImmunoResearch, 115-165-003) was used to identify BBB leakage in the brain sections. After rinsing with PBS, the preparations were incubated with appropriate secondary antibodies (Alexa Fluor 488, 555 or Cy3, Cy2 conjugated) for 60 min at room temperature. To visualize vessels, they were incubated with ‘Fluorescein’ or ‘Texas Red’ labeled Lectin (Vector Laboratories, Burlingame, CA). Control stainings with using only the secondary antibody were done to ensure if the detected signals were specific for the targets. Positive control stainings for RFC1 were done by labelling the sections involving choroid plexus, where the staining of RFC1 had already been established in literature (16). Colocalization studies were done from double immunohistochemical studies. The preparations were covered with Hoehst 33258 (Molecular Probes, ThermoFisher Scientific) that label cellular nuclei.

### The Isolation of Brain Microvessels

The brain was placed in ice cold HBSS and HEPES solution mixture in a petri dish. Under a stereomicroscope (Nikon SMZ745T) pineal gland, meninx and choroid plexus were removed, and the tissue was homogenized, then centrifugated at 2000 g for 10 min at 4 °C and supernatant was discarded. 18% 70.000 kDa Dextran was added on the pellet, centrifugated at 4.400 g for 15 min at 4 °C to separate non-vascular brain tissue. The pellet with vessels was resuspended and filtered through 100 µm and 40 µm filters to obtain microvessel fragments. The microvessels were transferred to adhesive slides for fixation or long-term preservation in -80 °C (17).

### Imaging and Image Analysis

The images of the stained brain preparations were obtained with a Leica TCS SP8 confocal laser scanning microscope (Leica, Wetzlar, Germany) with a diode laser 405, 638 and OPSL 488, 552 nm, with a X, Y, and Z-movement controller or a high-resolution PMT (Zeiss, Oberkochen, Germany) and HyD (Leica) detectors. All the analysis were done blindly to the groups. Images were either acquired in a single focal plane while keeping settings constant between experimental and control groups for analysis or Z-stack mode with 0.80 µm wide steps along the Z axis. For demonstration of RFC1 expression in brain sections and isolated microvessels, the images were taken in 2048x2048 pixels at 40X magnification (HCX APO L U-V-I 40x/0.80 water objective). For the analysis of immunostained ischemic brain sections, 3 regions of interest (ROIs) were manually inserted in each section, the infarct area (to represent the ischemic core), the adjacent cortical tissue (assumed to represent the periinfarct area) and the corresponding contralateral hemisphere were captured. Images were exported as 8-bit grayscale .tiff formats from Leica Application Suite X (LAS X; version 3.5.5.19976) and opened in ImageJ 1.52 version. ROIs were used to capture the fluorescence signal assigned to RFC1. These images were further processed with ImageJ (National Institutes of Health, Bethesda, MD, USA) to calculate the mean gray value in relative fluorescence unit (R.F.U.) to capture fluorescence intensity assigned to RFC1 staining. Since conditions of immunohistochemistry and confocal settings were kept constant no normalization of data is implemented and mean grey values were reported as arbitrary units.

### Western Blotting

Middle cerebral artery territories from the whole brains were cut in wedge shape from the brains of separate experimental group of mice. The ipsilateral adjacent brain tissue around the ischemic infarct was excised as the periinfarct area. The corresponding parts of the contralateral hemisphere were removed as controls. Infarct, periinfarct area, and contralateral parts were lysed in RIPA buffer containing protease and phosphatase inhibitor cocktail, homogenized on ice, centrifugated in 10000xg for 20 min at 4 °C. Protein concentration of the lysates was quantified using BCA protein assay kit. After loading proteins, 120 Voltage was applied on the gel for 1.5 h. By applying 120 mAmp for each membrane for 2 h in room temperature, they were transferred onto PVDF membrane. Membranes were blocked with 5% BSA in 0.1% TBS-Tween for 1 h at room temperature, then incubated with primary antibody against RFC1 (Sigma-Aldrich V44167) at +4 °C overnight. After washing with 0.1% TBS-Tween, they were incubated in goat anti-rabbit HRP conjugate (Invitrogen, 31460) solution in 2 h at room temperature. For detection, the membranes were treated with enhanced chemiluminescent (ECL) substrate (34094, Thermo Scientific™). Imaging was done by Kodak 4000M Image Station. β-tubulin III was used as an internal standard. The densitometric measurements were done by ImageJ 1.52 version (NIH, Bethesda, and Maryland), and were expressed as ratios to β-tubulin III values.

### Gene Silencing *in vivo* by Short Interfering RNA (siRNA)

To silence RFC1 gene, we pooled two of the specifically designed RFC1 (SLC19A1) targeted Accell siRNAs (Horizon Discovery, Waterbeach, United Kingdom). By choosing two separate siRNA targeted to different regions of RFC1 mRNA we aimed to enhance the potency and specificity. Accordingly, two separate non-targeting scrambled siRNAs were pooled as the control. The product codes and sequences are mentioned in our previous manuscript (8). We employed 2 different strategies when administering siRNA. Initially, RFC1-siRNA (50 µM) or scrambled-siRNA (50 µM) were intracerebroventricularly injected to mice via 27 G syringe (80308, Hamilton, U.S.A.) as a single dose, and their middle cerebral artery territories were extracted 48 h later. Subsequently, tissue RNA was isolated, and qRT-PCR was conducted. However, RFC1 mRNA silencing in the brain by intracerebroventricular RFC1-siRNA injection was unsuccessful. Hence, we targeted the medial rim of middle cerebral artery territory. We also doubled the concentration of siRNA to 100 µM and administered 1 µl of siRNA per injection, to avoid the overflow. Using the stereotaxic frame two burr holes targeting the middle cerebral artery territory was created (AP: 0/-1, ML: +2, DV: +2). The needle was then directed to the cortex in alignment with the curved cerebral surface. We used separate animals for RFC1-siRNA and scrambled-siRNA injections.

### Systemic Methotrexate (MTX) Administration

400 mg/kg of MTX (Koçak Farma, Ankara, Turkey) was injected to naive mice intraperitoneally (i.p.). The dose was chosen based on the literature showing observable changes within 24 h. Besides, the dose was chosen to be well below than maximally tolerated dose of MTX for systemic injections (18–20).

### Quantitative Reverse Transcriptase-Polymerase Chain Reaction (qRT-PCR)

siRNA-mediated RFC1 silencing was confirmed via the qRT-PCR method from the brains as described previously (8). The wedge-shaped fresh brain tissue from the injection site was extracted to isolate RNA via TRIzol Reagent (Invitrogen™) according to manufacturer’s manual. cDNAs were synthetized using High-Capacity Reverse Transcription cDNA Kit (Applied Biosystems). Subsequently, Applied Biosystems TaqMan® Gene Expression Assay including FAM™ dye labeled TaqMan® MGB probe and two unlabeled PCR primers for SLC19A1 gene (Assay ID#: Mm00446220_m1) were used. Reactions were performed in triplicates with the housekeeping gene TUBB4A (tubulin beta 4A class IVa, Assay ID#: Mm00726185_s1) as internal control. The difference in CT values (ΔCT) between the RFC1 (SLC19A1) gene and the housekeeping gene was then normalized to the corresponding ΔCT of the vehicle control (ΔΔCT) and expressed as fold expression (2−ΔΔCT) to assess the relative difference in mRNA expression.

### Magnetic Resonance Imaging (MRI) and Analysis

RFC1-siRNA or scrambled-siRNA injections were performed (n=22), and MCAo were performed 24-h later. Supplemental figure 1 depicts the experimental timeline of the MRI study. *In vivo* MRI were performed during occlusion, as previously described (21–24). In brief, a 7 Tesla horizontal-bore Bruker Avance I rodent imaging system (Bruker Biospin, Ettlingen, Germany), using a 72 mm inner diameter birdcage coil for transmission and a 15 mm diameter surface coil was used for reception. The *in vivo* MRI protocol comprised a diffusion-, perfusion-weighted, and spin-echo T2 weighted image axial sequences. During MRI, anesthesia was maintained with a mixture of air and 2% isoflurane (ISO-VET, Piramal Healthcare, Morpeth, UK). The animals were positioned on MRI-compatible mouse cradle. The respiratory rhythm and the body temperature were cautiously monitored by a pressure sensor linked to a monitoring system (ECG Trigger Unit HR V2.0, RAPID Biomedical, Rimpar, Germany). All MRI parameters are reported in Table 2. TOF-MR angiography was obtained to confirm successful occlusion of the MCA). Additionally, T2w images (anatomy), DWI (cytotoxic edema), and DSC-PWI (perfusion defect upon gadolinium injection) were acquired during the 90 min ischemia, before recanalization outside the magnet. The second MRI was performed at 24 h of the recanalization. To observe the BBB permeability, a T1-weighted dynamic contrast-enhanced (DCE) MRI sequence was used with 10 repetitions over 15 min and gadolinium administration after the third one. Mice were sacrificed via cardiac perfusion after the final MRI scan, and the brains underwent fixation, cryopreservation and cryosectioning as mentioned above for correlating immunohistochemistry.

**Table 2.**
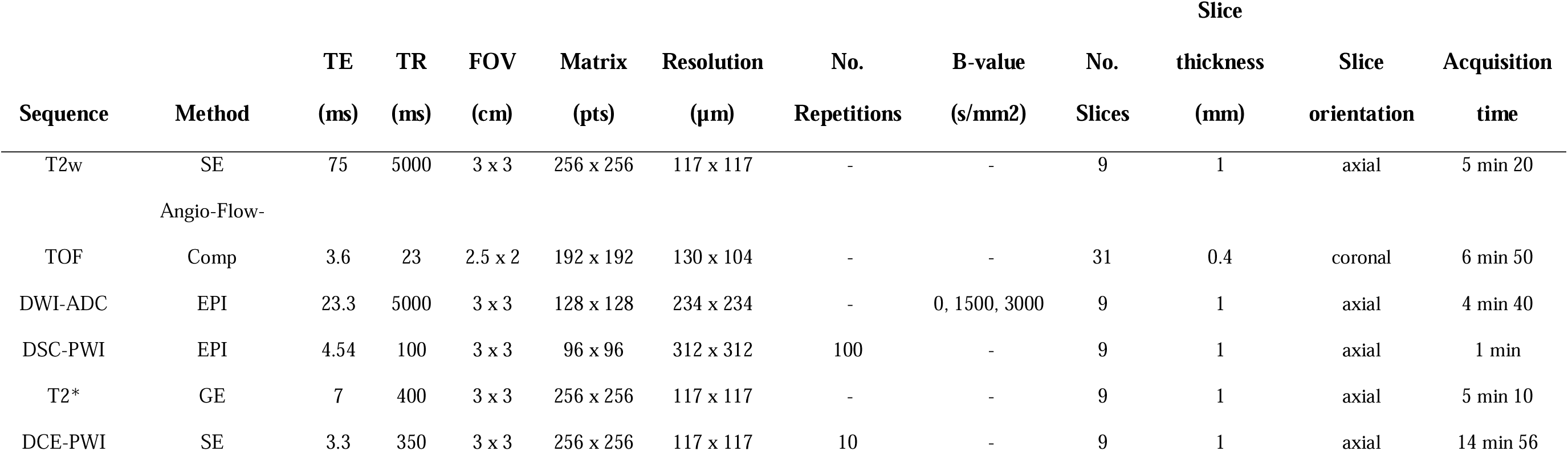
Parameters of the MRI acquisitions.

Raw data from Bruker were converted to Nifti format using MIPAV (http://mipav.cit.nih.gov/). AUC calculation was made as in Lemasson et al. (25). Manual delineation was performed using free hand tool in ImageJ 1.52 (National Institute of Health, USA imagej.nih.gox/ij/). Ipsilateral and contralateral hemispheres, along with the lesion of leakage of gadolinium were manually demarcated. Then, number of pixels multiplied by voxel size to obtain the volume (in mm^3^). The Gerriets correction for edema, as per the methodology proposed by Koch et al., was calculated (26). For comparing lesion changes over time, absolute infarct growth was calculated in cubic millimeters (mm³) by subtracting the final (T2w) lesion volume from the initial (ADC) lesion volume.

### Statistical analysis

A priori sample size estimation was performed based on our previous experiments for which infarct size was determined by histology in the same animal model. The expected mean infarct volume for 90 min of ischemia in Swiss albino mice is 44 mm^3^ (27), and the expected standard deviation is calculated as 10 in Swiss albino mice according to a literature meta-analysis (28). With a 30% decrease in infarct volume considered neuroprotective, the effect size is 1.26. Using G-Power 3.1.9.4 (difference between two independent means, alpha error 0.05, power 0.8), the sample size for the experimental and control groups was found to be n = 9 animals per group.

Data were analyzed using IBM SPSS 23 statistical analysis program. Normally distributed data were tested using a T test or analysis of variance (ANOVA) followed by post-hoc Fisher’s LSD. Non-normally distributed data such were compared using the Mann-Whitney U test (for two groups), and Kruskal Wallis (for more than two groups). P values were given in respective figure legends, p ≤ 0.05 was considered significant. Results were expressed as mean ± standard error of the mean (S.E.M.).

## RESULTS

### Microvessels of the brain express RFC1 profusely

RFC1 protein expression was evidenced in mouse brain sections and isolated microvessels. Notably, RFC1 was prominently observed in α-SMA positive capillaries (Figure 1A, arrows) and in capillaries with < 5 µm diameter (Figure 1A, arrowheads). Staining with astrocytic endfeet marker AQP4 revealed that RFC1 was not expressed in astrocytic end feet but localized adjacent to it (Figure 1B). Since thin brain sections usually do not allow visualization of entire microvessels including bifurcations, it is essential to use thick sections, but RFC1 antibody needs to be used at high concentrations and for long incubation periods to achieve adequate staining. Therefore, we isolated cerebral capillaries and stained RFC1 concomitantly with tomato lectin that label microvessels or with mural cell markers (PDGFR-β, CD13). Lectin labelled glycocalyx around endothelial cells and pericytes which colocalized with RFC1 staining. RFC1 also colocalized with pericyte markers (PDGFR-β, CD13) indicating that RFC1 is expressed in pericytes as well (Figure 1C-E).

**Figure 1.**
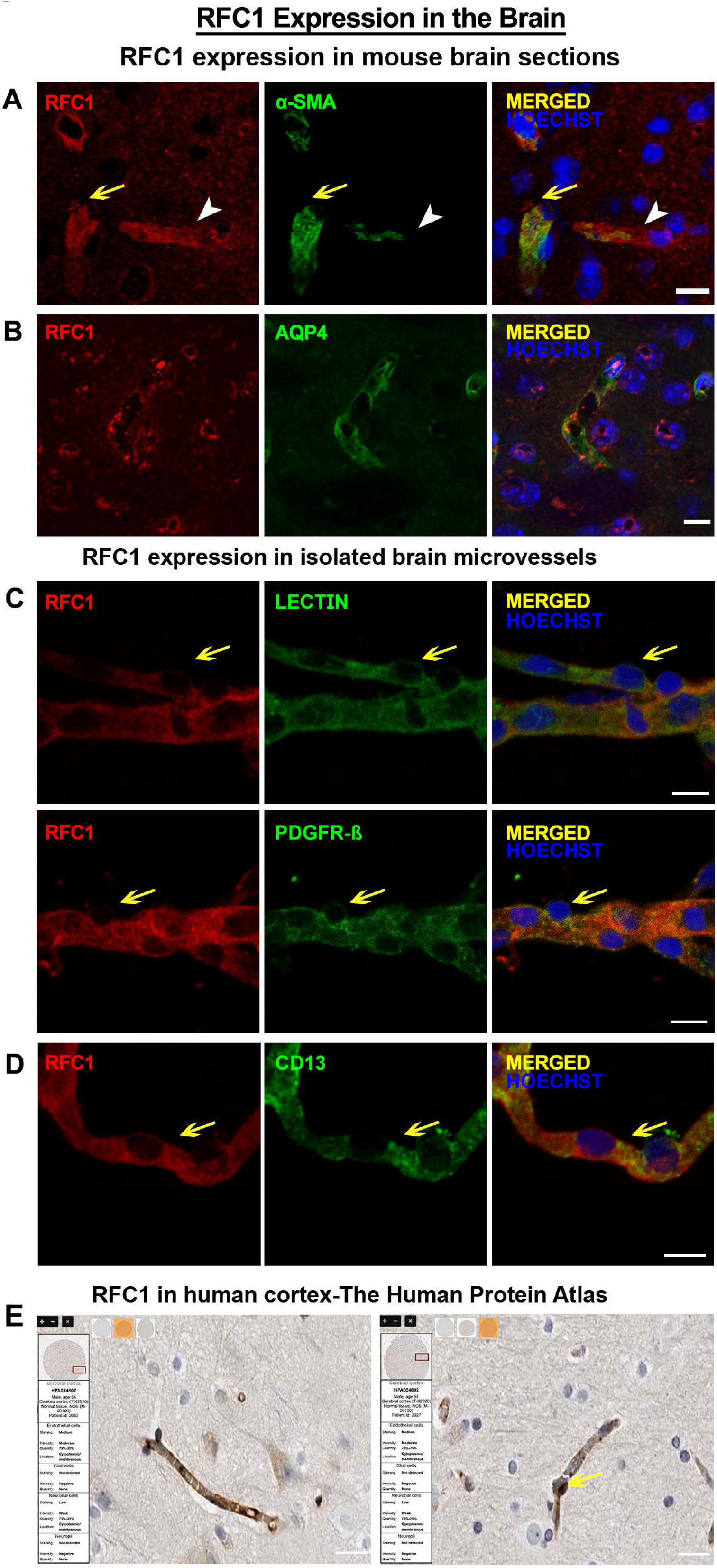
RFC1 protein is expressed in the brain microvessels. A) PFA fixated 20 µm-thick brain sections of naive Swiss Albino mice was labelled with anti-RFC1 antibody (red), and α-SMA (green). RFC1 is expressed in α-SMA positive precapillary arterioles (arrows) as well as capillaries (arrowheads). B) RFC1 is not expressed in AQP4 positive astrocytic perivascular endfeet rather the RFC1 immunoreactivity is observed in spatially distinct regions. C) Isolated brain microvessel (< 9 μm diameter) preparations were immunohistochemically labelled with anti-RFC1 antibody (red), and colocalized with the vessel marker Lectin (green). The bump-on-a log pericyte shape can be observed as RFC1 positive (yellow arrows). D) The common pericyte marker PDGFR-β (green) colocalized with RFC1. Yellow arrow depicts a pericyte body appearing as a bump-on-a log on the microvessel. E) CD13 (green) also colocalized with RFC1. Nuclei were labeled with Hoechst 33258 (blue). Scale bars: 10 μm. F) RFC1 labeling in the post-mortem human cortex also showed that RFC1 is expressed in the endothelial cells and pericytes (yellow arrow) (www.proteinatlas.org). Scale bars: 20 μm

We also compared our results to RFC1 stainings in post-mortem human brain tissue in Human Protein Atlas which confirmed that RFC1 protein is expressed in the brain endothelial cells in both human and mouse microvessels (29). However, the atlas did not mention pericytes although upon careful examination anti-RFC1 antibody visibly labeled the pericytes, which can be distinguished by its distinct morphology, that is, -protruding ovoid shaped **cell bodies** with the classic **bump-on-a-log** appearance-(Figure 1F, arrow).

### RFC1 levels were downregulated in the ischemic core but upregulated in the periinfarct area 24-h after ischemia

To test if RFC1 levels were temporally and spatially affected by ischemia, we performed intraluminal transient focal cerebral ischemia model with different recanalization time-points (1 h, 24 h, 48 h) following 90 min ischemia. We studied the protein levels in the infarct, periinfarct, contralateral areas via immunohistochemical studies and Western blotting.

Wedge resections from these defined regions revealed that RFC1 protein levels decreased in the ischemic core promptly (71% decrease 1 h after ischemia compared to the average level of naïve brains). This decrease persisted compared to contralateral also in other time-points (38% decrease at 24-h and 26% at 48 h) (Figure 2A). The average values in the contralateral side were stable between time-points, and similar to naïve values. We observed that RFC1 expression in periinfarct are slightly elevated, with 36% increase at 24 h, and 32% increase at 48 h after ischemia compared to contralateral although this did not reach to statistical significance at 24 h (Figure 2B). We characterized RFC1 immunoreactivity in ischemic brains (24 h after ischemia) which was apparent and diffuse throughout the contralateral brain hemisphere. However, the core ischemic area stained weakly for RFC1 compared to periinfarct and contralateral areas (Figure 2C, D). Spatially controlled quantitative analyses of fluorescence intensity in ischemic brain sections immunostained with anti-RFC1 antibody, revealed that fluorescence intensity was increased in the periinfarct region in the 24 h recanalization group (Figure 2E).

**Figure 2.**
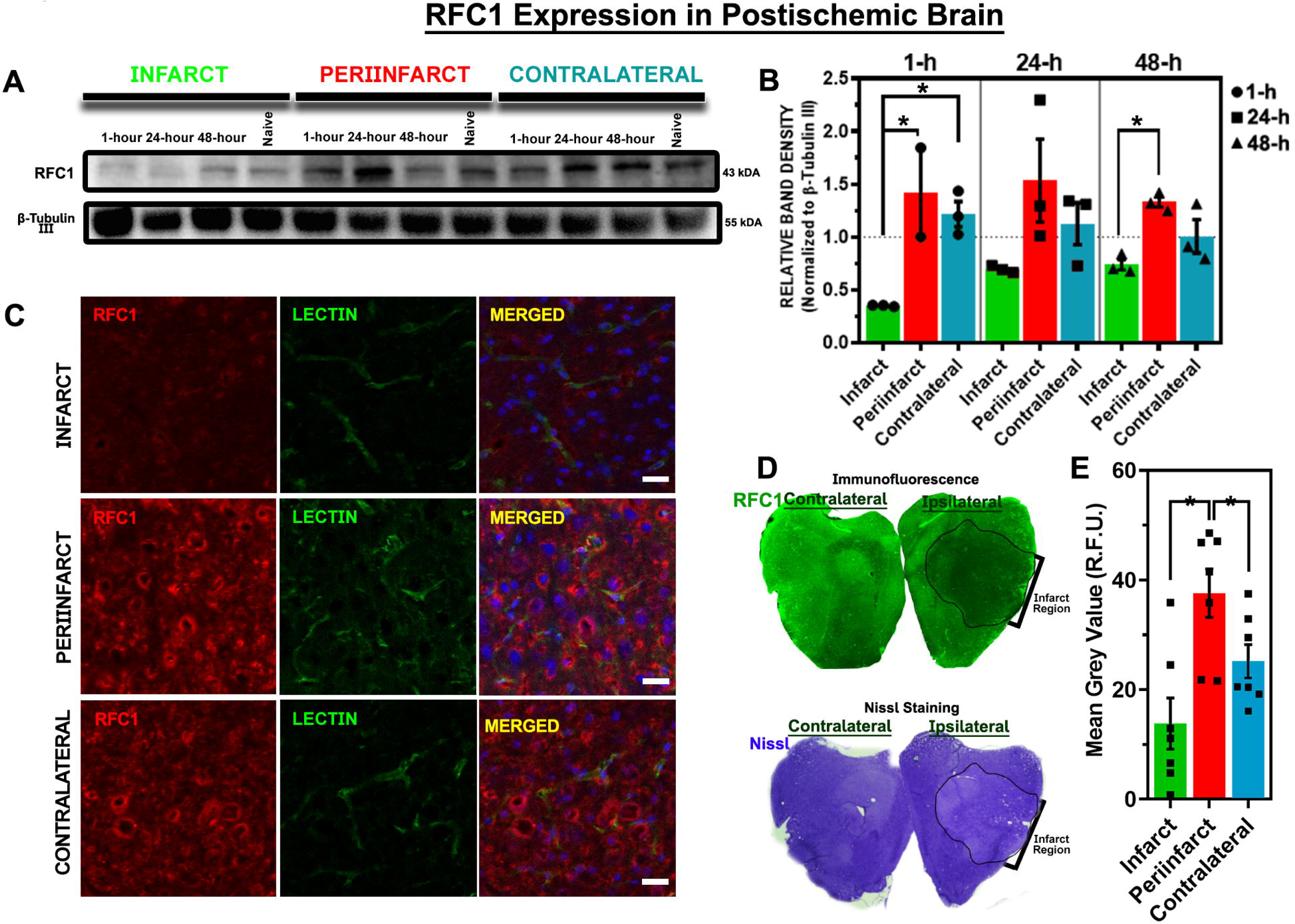
RFC1 levels are dynamic throughout the pathophysiology of ischemic stroke. Representative Western blot showing RFC1 protein expression in the core, periinfarct, and contralateral tissues from naïve and ischemic mice brains following 90 min ischemia, with recanalization at 1 h, 24 h, or 48 h. The band at ∼43 kDa corresponds to RFC1 protein, showing increased expression in the periinfarct region of the ischemic hemisphere. B) Densitometric quantification of Western blotting was normalized to the average of naïve brains (n=3), and to β-Tubulin III as the loading control. Results are expressed as relative optical density and represented as mean ± S.E.M. The average optical density of naïve brains (n=3) is represented as the horizontal dashed line in the graph. RFC1 protein expression significantly decreased in the core region at all time points and increased in the periinfarct region at 1 h, 24 h, and 48 h on average. One-way ANOVA followed by post-hoc uncorrected Fisher’s LSD was used for comparisons. For 1 h: infarct vs. periinfarct (p=0.0107) infarct vs. contralateral (p=0.0158), periinfarct vs. contralateral (p=0.4812). For 24 h: infarct vs. periinfarct (p=0.0581), infarct vs. contralateral (p=0.2754), periinfarct vs. contralateral (p=0.2989). For 48 h: infarct vs. periinfarct (p=0.0057), infarct vs. contralateral (p=0.1060), periinfarct vs. contralateral (p=0.0613). C) Representative confocal images at the 24 h recanalization time-point showed decreased RFC1 immunostaining in the infarct region and increased RFC1 in the periinfarct region compared to the contralateral. D) Nissl staining done on the adjacent section to confirm the infarct region, and delineate between core, periinfarct and contralateral regions. E) Mean Grey Value (R.F.U.) as an indicator of fluorescence intensity was measured from ROI’s manually placed in core, periinfarct, and contralateral regions. Mean grey value of periinfarct region found higher than core and periinfarct regions at 24 h time-point. One-way ANOVA followed by post-hoc uncorrected Fisher’s LSD comparisons: infarct vs. periinfarct (p=0.0107), infarct vs. contralateral (p=0.0158), periinfarct vs. contralateral (p=0.4812). S.E.M.: Standard Errors of the Mean; R.F.U.: Relative Fluorescence Unit. Scale bar= 20 µm.

### RFC1 knockdown aggravated BBB breakdown in periinfarct area but did not affect infarct size 24-h after ischemic stroke

Unnoticed for a long time in cerebral microvessels, the role of RFC1 in the brain via modification of RFC1 protein *in vivo* was not studied. To date, there is no commercially available RFC1 transgenic animal. Furthermore, knocking-out RFC1 in mice was incompatible with life (30). Therefore, we opted to modify the RFC1 levels using siRNA or a pharmacological agent. Based on our success with modification of RFC1 levels in mouse retina, we decided to utilize RFC1 targeted Accell siRNAs (RFC1-siRNA) or corresponding scrambled siRNAs as fast and efficient tools to modify RFC1 levels in the brain (8). For pharmacological approach, we utilized a non-covalent (competitive) inhibitor of RFC1, MTX, to modify the RFC1 levels since there are no specific pharmacological inhibitors available.

First, we delivered a pool of 50 µM RFC1 targeted Accell siRNAs (RFC1-siRNA) or scrambled siRNAs intracerebroventricularly. We performed qRT-PCR from whole brain homogenates resulting in no change in RFC1 levels 48 h later (data not shown). Then, we doubled the dose to 100 µM and shortened the incubation period to 24 h and injected the pool of siRNA intracortically. Performing qRT-PCR from the injection site, we observed a regional RFC1 knock down via qRT-PCR (Fig. 3A) and immunohistochemistry (Fig. 3B, C). RFC1 mRNA levels decreased by 30% at the site of injection (Figure 3A, p=0.028).

**Figure 3.**
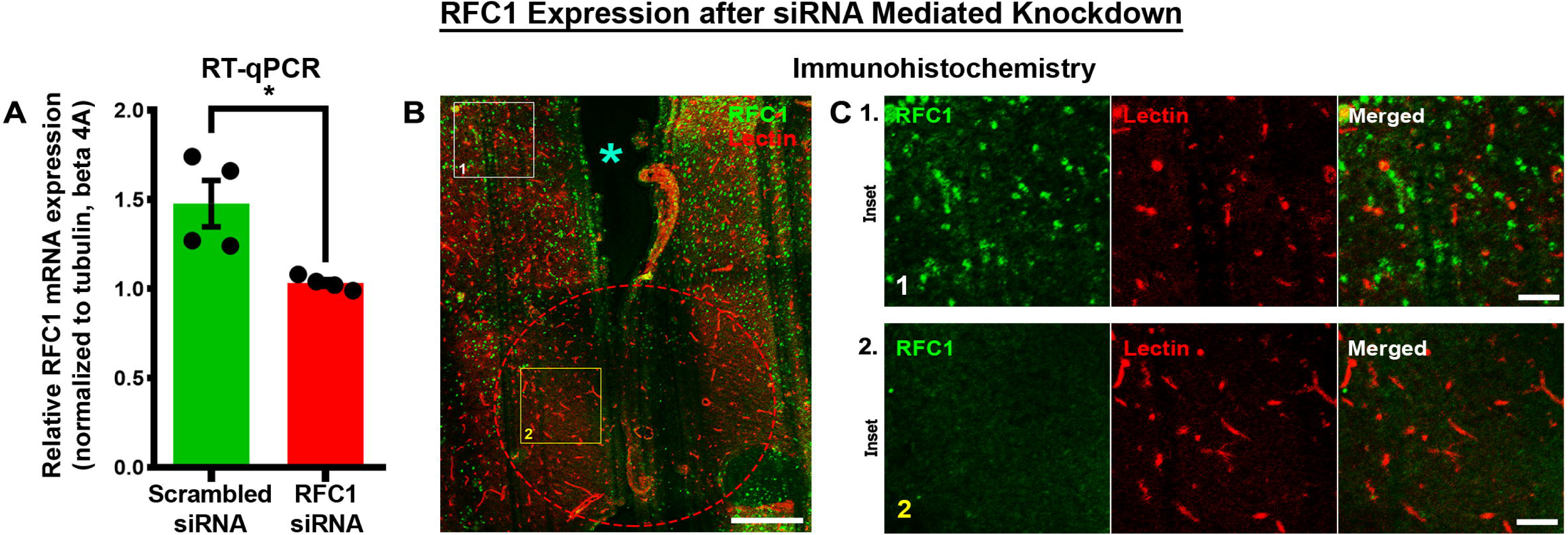
Intracortical injections of RFC1-siRNA resulted in local RFC1 knockdown. A) The bar graph shows that intracortical RFC1-siRNA delivery reduced RFC1 mRNA levels by 30.1% when compared to scrambled-siRNA delivered mice brains (*p= 0.028). B) The wedge-shaped injection site is shown (cyan asterisk) in a representative brain section. RFC1 immunoreactivity (green) was decreased locally in the affected region demarcated by a circular red dashed line. Scale bar= 250 µm. C) Insets depict the local RFC1 knock downed. 1) At higher magnification, positive RFC1 staining was observed outside of the injection site. 2) However, within the injection site, RFC1 immunoreactivity disappeared. Scale bar= 50 µm.

Next, we delivered the pool of RFC1-siRNA (n=12) or scrambled siRNA (n=10) to the medial rim of the middle cerebral artery territory, a potential periinfarct area of cortex. We then performed intraluminal MCAo (n=21) or sham surgery (n=1) 24 h later in RFC1-siRNA injected mice. After the occlusion of MCA, per-occlusion multimodal MRI confirmed the occlusion (angiography) and highlighted a typical cortico-striatal acute infarct (diffusion/perfusion imaging). Animals were excluded from RFC1-siRNA (n=1) and scrambled-siRNA (n=1) groups due to surgical failure (incomplete occlusion), at this point. Several animals from RFC1-siRNA (n=5) and scrambled-siRNA (n=3) group died during the night possibly due to lethality of MCAo. Next day, we performed the modified Bederson scoring on surviving animals from scrambled-siRNA (n=6), RFC1-siRNA (n=6) and sham (n=1) groups. The median and interquartile ranges of the Bederson Scoring for the RFC1-siRNA group and scrambled siRNA group were 3.5 [2.5–4.25]) and 3.0 ([2.5–4.0]) respectively, implying the severity of stroke between the groups was similar. Additionally, one more animal from the RFC1-siRNA group, which had a Bederson score of 5, had died before the second set of MRI scans could be conducted. A second session of MRI was performed 24 h after recanalization, so as to measure the final lesion (T2w), and to evaluate BBB integrity (DCE-MRI) (Fig. 4A-B).

**Figure 4.**
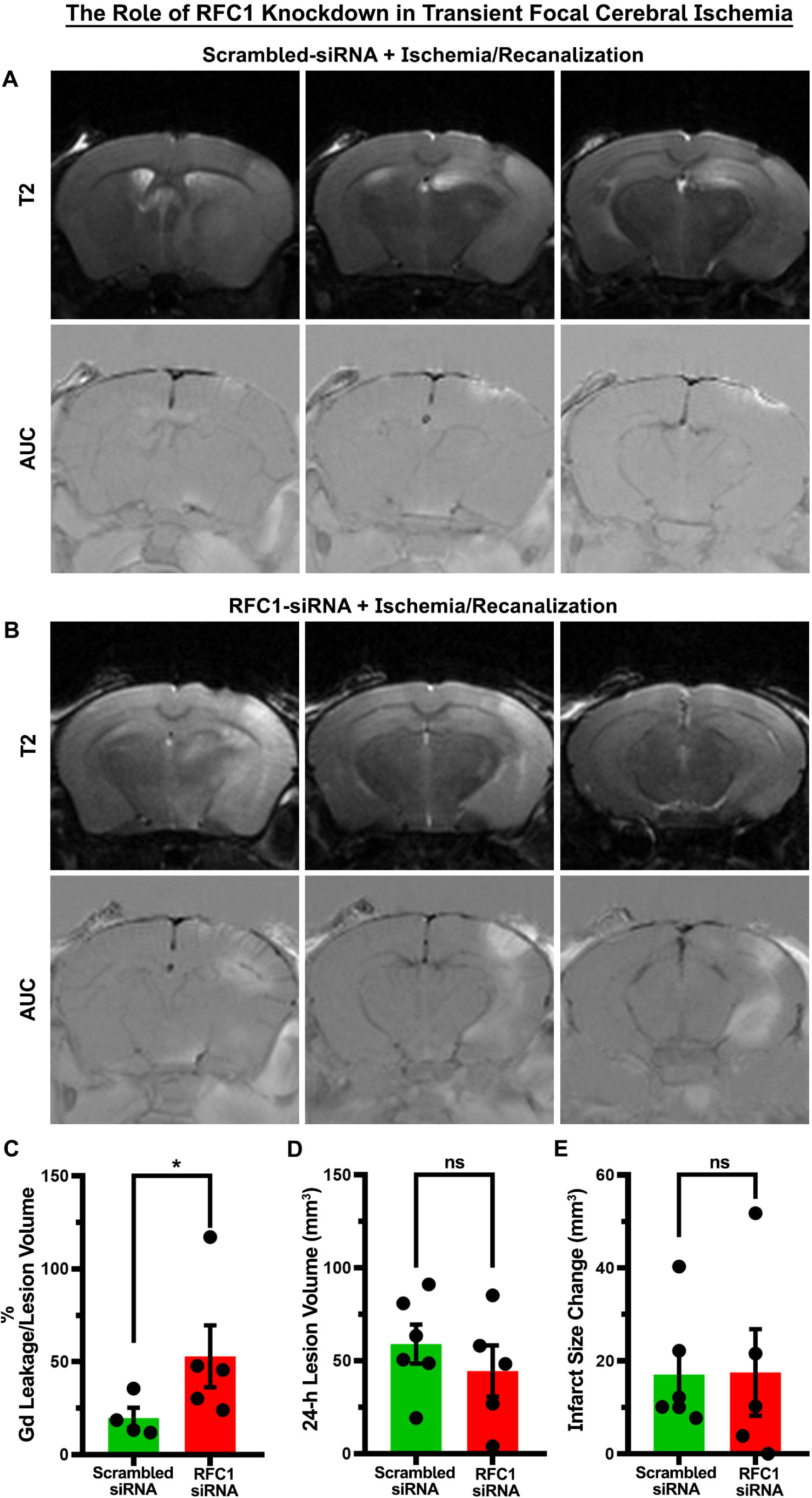
RFC1-siRNA administration before ischemia increased Gadolinium leakage from the BBB at 24. **h.** RFC1-siRNA or scrambled-siRNA were administered intracortically before ischemia/recanalization. 24 h after recanalization, T2 weighed as well as dynamic contrast enhanced T1 weighed MRI sequences were obtained. BBB dysfunction was evaluated by *in vivo* Gd leakage in AUC maps obtained from dynamic contrast enhanced T1 weighed images. The panels in A and B demonstrate 4 slices of T2 weighed MRI images and matched AUC maps of either a scrambled-siRNA or RFC1-siRNA administered ischemic brains after 24 h in recanalization. A) Wide cortical and striatal hyperintensities in the MCA territory in the upper panel indicates a successful ischemia induction in scrambled-siRNA injected brain. Despite this, minimal Gd leakage is observed in the injection site as indicated by bright white regions in the AUC maps. B) Other mice which received RFC1-siRNA showed a similarly comprehensive infarct in cortex and striatum. However, as seen in the representative MRI scans of one mouse, the matched AUC maps RFC1-siRNA injection to the cortex before ischemia caused substantially more Gd leakage from the BBB than the scrambled-siRNA injected ones. C) Since BBB disruption was expected in ischemia even without RFC1 knockdown, the total leakage detected in cortex was divided by cortical infarct volume for normalization. The percent of the cortical leakage volume in ratio to cortical infarct volume was significantly higher in RFC1 knocked down ischemic brains (n=5) compared to scrambled siRNA injected ischemic brains (n=4; p=0.03). D) However, 24 h cortical ischemic lesion volume calculated from T2 weighted images were not different between groups (n=6 for scrambled-siRNA and n=5 for RFC1-siRNA). E) The change in the infarct size over 24 h which was calculated by subtracting acute lesion size measured in DWI sequence from 24 h lesion size measured in T2 weighed sequence was not statistically significant between groups.

The acute cortical infarct volumes, 24 h cortical lesion volumes, and the change in the infarct size or cortical edema over 24 h did not exhibit statistical differences between groups (Fig. 4D-E). RFC1-siRNA treated ischemic brains demonstrated broader Gd leakage compared to scrambled-siRNA treated ones (Figure 4A, B). Moreover, BBB degrading caused by RFC1 knockdown was observed both in RFC1-siRNA injected sham animal or when RFC1-siRNA is injected in deep brain structures (Supplemental Figure 2). Since ischemia is known to impair BBB integrity and lead to Gd leakage without any genetic interventions, we normalized the cortical Gd leakage volume (mm^3^) to the cortical infarct volume (mm^3^). The resulting percentage of the cortical Gd leakage volume in ratio to cortical infarct volume was significantly higher in RFC1-knocked down ischemic brains compared to scrambled siRNA injected ischemic ones (Figure 4C, p=0.03).

As a complementary, pharmacological approach, we systemically administered MTX to modify RFC1 levels. MTX, a high affinity substate of RFC1 and competitive inhibitor of folate transport via RFC1 has been shown to increase RFC1 levels as early as 15 min whereas it decreased RFC1 levels in 72 hours in *in vitro* studies (31,32). Therefore, we decided to examine brain RFC1 levels 1, 24 and 48 h after animals that had been treated with a single high dose i.p. MTX injection (400 mg/kg). Immunofluorescent labelling and confocal imaging of MCA region, as well as Western blotting of whole brain homogenates revealed that systemic MTX increased the level of RFC1 in the brain after 1 h. In contrast, RFC1 levels decreased beyond naïve levels 48 h after MTX (Supplemental Figure 3A). When we investigated the BBB integrity in those brains, we did not observe any IgG in the parenchyma at 1 h or 24 h later. However, we observed IgG leakage 48 h after high dose systemic injection of MTX, coinciding with the timeline when MTX suppressed RFC1 levels (Supplemental Figure 3B).

## DISCUSSION

RFC1 expression, among other folate transporters in the brain, has been shown to be expressed in neurons, endothelial cells, astrocytes, and microglia in the brain (16). However, this study disregarded the mural cells (vascular smooth muscle cells and pericytes) and failed to address RFC1 expression in brain mural cells even though it was proposed as one of the only three common transcripts in all mural cell types in the most recent brain mural cell transcriptome atlas. While RFC1 protein in microvessels was not shown, and its potential role in the adult brain was overlooked before recent research conducted on human brain autopsies nominated RFC1 as one of the pericyte markers in paraffin embedded human brain sections (33).

In this study, we shed light on these facets of RFC1: 1) we demonstrated the expression of RFC1 in cerebral pericytes and endothelial cells, 2) we identified acute ischemia/recanalization related changes of RFC1 and 3) subsequent functional impact of its modification on BBB integrity.

We illustrated that RFC1 was colocalized with known mural cell markers (α-SMA, PDGFR-β and CD13) in the microvessels of the mouse brain. The reason that RFC1 at the protein level had gone unrecognized in pericytes so far might have several explanations. First, in the absence of RFC1 antibody, early studies used anti-serum to determine RFC1 immunohistochemically. Thus, lack of validated commercial antibodies prevented researchers from studying the transporter immunohistochemically. Second, although RFC1 was determined in brain sections of mice, rats, and post-mortem humans, the immunopositivity of RFC1 in the pericytes was unnoticed since pericytes was largely neglected by most of the neuroscientists. Hence, most studies evaluating the role of RFC1 in microvessels concentrated on merely the endothelial cells, but not the neighboring pericytes (34–37). Accordingly, a recent immunohistochemical characterization study which focused on the localization of different type of folate transporters in mouse brain successfully demonstrated RFC1 expression in brain barriers (including BBB and BCSFB); however, this study also omitted the pericytes among other cerebral cell types (16). Finally, a recent paper confirmed our findings by immunohistochemically labeling formalin-fixed paraffin-embedded (FFPE) human brain tissues showing human brain pericytes also express RFC1. In that study, the authors also discussed that RFC1 might be potential pericyte marker in humans, which was supported by single-cell RNA-seq analyses and immunoblots (33). Our findings along with this recent human histopathological study, confirmed the results of a meta-analysis that highlighted the RFC1 gene as one of the only three consistently detected genes in brain pericytes out of 1180 other genes (38).

To understand if RFC1 has a role in acute ischemic stroke pathophysiology, we induced 90 min ischemia followed by 1 h, 24 h, or 48 h recanalization, and detected if RFC1 protein varied in relation to brain regions and timepoints. We found that RFC1 protein expression was suppressed in the core region at these three time-points. Additionally, this effect was not due to antigen hindering effect of ischemia since Western blots were also confirmed our findings. Moreover, RFC1 was detected higher in periinfarct region compared to contralateral. Since there is no other comparable specific study focusing on RFC1 levels in stroke, we turned back to -omic studies examining broad spectrum of mRNAs or proteins in various forms of ischemic modalities. We determined that RFC1 has not been included in proteomics studies consisting either permanent (39) or transient (40–42) ischemic stroke models inquiring acute (40–42) or chronic time points (39,43). Transcriptomic studies investigating mRNA levels in acute or subacute forms of transient MCAo (60 min occlusion followed by from 24 h to 7 d of recanalization) reported no alteration of RFC1 in ischemic microvessels (44), astrocytes (45) or cerebellar cortex (46). On the other hand one study reported that RFC1 is induced in microglia in 14 days after permanent ischemic stroke in young mice, despite being statistically insignificant (47). In another study, no variation was detected in RFC1 expression in ischemic endothelial cells during the acute (24 h) or subacute (72 h) phases. However, the chronic (1 month) timeframe revealed an upregulation (48).

The genetic elimination of RFC1 is lethal and no conditional knockout animals were ever used in the literature before; therefore, we opted for siRNA technology to suppress RFC1 levels in the brain (49). Earlier studies using RFC1-siRNA were conducted in primary cultures of human differentiated adipocytes, and in rat choroidal epithelial Z310 cells; however, those studies were limited to *in vitro* (50,51). Hence, our previous paper was the initial study achieving the knockdown RFC1 *in vivo* (8). Due to inconclusive knockdown efficiency of RFC1-siRNA when administered intracerebroventricularly, the siRNA was administered into cortex. This resulted in a local siRNA-mediated knockdown of RFC1, disrupting BBB integrity, and causing extravasation of endogenous IgG, a phenomenon has been showed in the retina in our previous study.

To further investigate the interrelationship with ischemic stroke and RFC1 expression, we injected RFC1-siRNA and scrambled-siRNA into the anticipated periinfarct cortex of mice and induced ischemic stroke in the following day. The results were monitored 4 h after the stroke via 7T MRI, the gold standard non-invasive method for clinical diagnosis and follow-up for ischemic stroke treatment. The most prominent outcome was the increase of the Gd leakage in the brain treated with RFC1-siRNA. After adjustment for lesion size, brains injected with RFC1-siRNA still showed an increased BBB disruption. However, in the H0 infarct (derived from per-occlusion DWI) and H24 lesion (measured on T2w on the following day) were not different among groups. While an increase in infarct size could be expected after BBB impairment due to RFC1 reduction, several explanations can be put forward. First, we might have not been able observe the possible infarct size change due to locally restricted effect of siRNA in the periinfarct cortex. The local impact of intracortical siRNA when contrasted with a large cortical infarct size, might have hindered our ability to determine the effect of RFC1 reduction on infarct size. Second, compensatory mechanisms might have mitigated the effect of RFC1 reduction on ischemic infarct evolution. Third, timing of the assessment, limited to 24h in this study, might have also hindered the potential contribution of RFC1 knockdown to infarct size development.

Our experimental timeline with MRI required prolonged anesthesia exposure, combined with the intraluminal filament technique, which has high mortality. As a result, 6 RFC1-siRNA and 3 scrambled-siRNA injected ischemic mice died during the protocol. We could not observe hemorrhage after 24 hours in this cohort of animals. However, given the prominent effect on BBB, we still consider RFC1 as a candidate which might play a role in hemorrhagic transformation. Future studies, using other models of stroke such as the thrombotic model or FeCl_3_ induced distal MCAo model, are needed to better characterize the functional consequence of this increased permeability of the BBB after RFC1 downregulation (52,53).

The effects of MTX, a drug that functions as a folate analogue, and a non-covalent (competitive) inhibitor of RFC1 were investigated for its impact on RFC1 levels. The results obtained were significant as they also provide a timeline for the effects of single dose systemic MTX. It was observed that the level of RFC1 in the whole brain increased in the first hour following administration of a single dose systemic MTX administration. This increase might have been because MTX, a competitive inhibitor of folate transport via RFC1, reduced the intracellular transport of folate through RFC1, thus the cells increased RFC1 levels to meet the constant folate need (54). We determined that the BBB was compromised 48 h after high-dose MTX administration. Surprisingly, this coincided with the time point when RFC1 levels were suppressed following MTX intervention. We acknowledge that this synchronicity cannot be solely attributed to the effect of MTX on reduction of the RFC1 levels, since MTX is not a specific substrate of RFC1 and also competitively inhibits the activity of folate cycle enzymes and synthesis of purine and pyrimidine required for DNA and RNA production. However, coupled with our results from RFC1-siRNA studies, we consider that suppression of RFC1, as well as the inhibition of the folate cycle, might result in failure to utilize folate, leading to the disruption of BBB (55).

These findings support a recent clinical study investigating the association between single nucleotide polymorphisms (SNPs) of RFC1 gene and ischemic stroke, where some RFC1 polymorphisms were found more frequently in small arterial occlusions and silent brain infarctions (6). Furthermore, these SNPs were not found to be directly linked to life-time low folate levels or hyperhomocyteinemia, both of which also create susceptibility to stroke. Additionally, there is a lack of information regarding how these SNPs may alter RFC1 in response to various stress factors and RFC1 modifiers (e.g., NRF-1, HIF1α, and NO), which may lead to dysfunctional or reduced RFC1.

Here, we propose that proof of concept study may provide insight to the fact that RFC1 dysfunction might compromise the BBB function, hence might worsen ischemic stroke. More comprehensive studies breaking down the role of RFC1 is required to understand the role of it in ischemia/reperfusion or its impact on the BBB.

## ABBREVIATIONS

BBB: blood-brain barrier
BRB: blood-retina barrier
CD13: Aminopeptidase N
IgG: Immunoglobulin G
mRNA: messenger RNA
MTX: Methotrexate
PBS: phosphate buffered saline
PDGFR-β: platelet derived growth factor receptor beta
PFA: paraformaldehyde
qRT-PCR: Quantitative Reverse Transcriptase-Polymerase Chain Reaction
RBF: Relative blood flow
RFC1: Reduced Folate Carrier 1
RIPA: Radioimmunoprecipitation assay
siRNA: short interfering RNA
SLC19a1: solute carrier family 19 (folate transporter), member 1
SNPs: single nucleotide polymorphisms

## DECLARATIONS

### DATA AVAILABILITY

There are no restrictions on data availability. Source data are provided with this paper. Supplementary information is available for this paper. Correspondence and requests for materials should be addressed to MY or GG.

### COMPETING INTERESTS

The authors declare that they have no competing interests.

### FUNDING

This project received a bilateral France-Turkey grant (BOSPHORUS program, n° 46602VB). This research was supported by The Scientific and Technological Research Institution of Turkey (TÜBİTAK; Grant No: 120N690) and Hacettepe University Scientific Research Coordination Unit (Project No: TDK-2020-18590).

### AUTHORS’ CONTRIBUTIONS

GG contributed to the study design, performed surgeries and experiments, obtained tissues, performed immunohistochemistry, collected images and data, analyzed the data, prepared figures, wrote the original manuscript and contributed to editing. DB performed experiments, contributed to image collection, and obtaining tissues. CL performed surgeries. NB contributed to design and execution of Western Blottings and collected images. MSB contributed to design and execution of qRT-PCR experiments and performed related analysis. RB contributed to the in vivo MRI experiments and data analysis. HK, contributed to the methodology and provided resources. MW contributed to the application of MRI studies, contributed to the funding acquisition and editing of the paper. FC contributed to the funding acquisition, design and application of MRI studies, performed surgeries and contributed to writing of the paper. MY acquired funds and administered the project, coordinated, and supervised the work, and contributed to figures and reviewed and edited the paper. All authors read and approved the final manuscript and provided minor modifications.

## ACKNOWLEDGMENTS

We thank Dr. Canan Cakir Aktas (The Institute of Neurological Sciences and Psychiatry, Hacettepe University, Ankara, Turkey) for assisting the financial management of the project, Dr. Buket Nebiye Demir (The Institute of Neurological Sciences and Psychiatry, Hacettepe University, Ankara, Turkey) for technical advice on in vivo procedures, Dr. Cetin Demir (Faculty of Medicine, Department of Pediatrics, Division of Pediatric Oncology, Drug Resistance Laboratory, Hacettepe University, Ankara, Turkey) for his assistance in the execution of qRT-PCR experiments, and our laboratory technician Mesut Firat (The Institute of Neurological Sciences and Psychiatry, Hacettepe University, Ankara, Turkey) for providing technical assistance throughout project. We are grateful to our undergraduate researchers Derin Nalcakan, Neslihan Nisa Gecici, Mustafa Guvercin (Hacettepe University Medical School), Asya Unal (Ankara University Medical School) who were officially added as interns to the project with “TÜBİTAK-Intern Researcher Scholarship Programme”, for their contribution to immunohistochemistry and data analysis.

## SUPPLEMENTAL FIGURE LEGENDS

**Supplemental Figure 1. Experimental timeline.** Mice were intracortically injected with 100 μMLJof siRNA at day 0. On the 1^st^ day animals were catheterized for administration of intravenous Gd during MRI. Subsequently, MCAo was induced, and immediate MRI was obtained. After 90 min of ischemia, mice underwent recanalization. On the 2^nd^ day, they were re-catheterized for Gd injection, and the second MRIs were performed. Afterwards, mice were sacrificed for post-mortem immunohistochemical studies and analyses.

**SupplementalFigure 2. siRNA mediated RFC1 knockdown led to BBB disruption and endogenous IgG extravasation.** A) The T1-contrasted MRI was obtained 24 h after the sham surgery following cortical RFC1-siRNA injection. The Gd leakage is seen as regional hyperintensity as shown by an arrow. B) After *in vivo* MRI, we sacrificed the same mice via cardiac perfusion and cryosectioned the brain to label with vessel marker Lectin (cyan) and Cy3 goat anti-mouse antibody (yellow) to visualize the extravasated endogenous mouse IgG (155 kDa). Cortical injection site is marked with the asterisk. RFC1-siRNA mediated knockdown led to significant IgG extravasation from the microvessels. Scale bar: 50 μm. C) Moreover, we investigated whether RFC1 knockdown resulted in BBB disruption when administered to the regions other than cortex. Intrathalamic siRNA injection resulted in IgG leakage along with the injection trace. The Hamilton injector was introduced 1700 μm deep into the brain in (ML:1.8, AP: -1.4, DV: 3.2). Syringe was retracted while slowly administering the siRNA over minimum of 5 minutes. Accordingly, thalamic nuclei (tn), fimbria (fi), and sensorial cortex (sc) along the trace showed IgG leakage. Scale bar=500 μm.

**Supplemental Figure 3. The effect of a single dose systemic MTX on RFC1 levels and the BBB integrity.** After a single dose MTX (i.p.), mice sacrificed at 1, 24 and 48 h. Immunofluorescent images and Western blotting of the brains of each mouse suggested that single dose of systemic MTX increased RFC1 protein levels at 1 h, and RFC1 protein was at the lowest at 48 h. Extravasation endogenous mouse IgG (155 kDa) was investigated at the same time points via immunohistochemistry. While 1 h and 24 h after MTX administration revealed no compromise in the BBB, 48 h time point showed IgG leakage in the parenchyma. Scale bar= 50 μm. Inset scale bar=10 μm.

## Notes

### Competing Interest Statement

The authors have declared no competing interest.

