## Supplementary figures and images for "Reduced Folate Carrier 1 (RFC1/Slc19a1) Suppression Exacerbates Blood-Brain Barrier Breakdown in Experimental Ischemic Stroke in Adult Mice"

### Supplemental Fig 2_No layers.tif

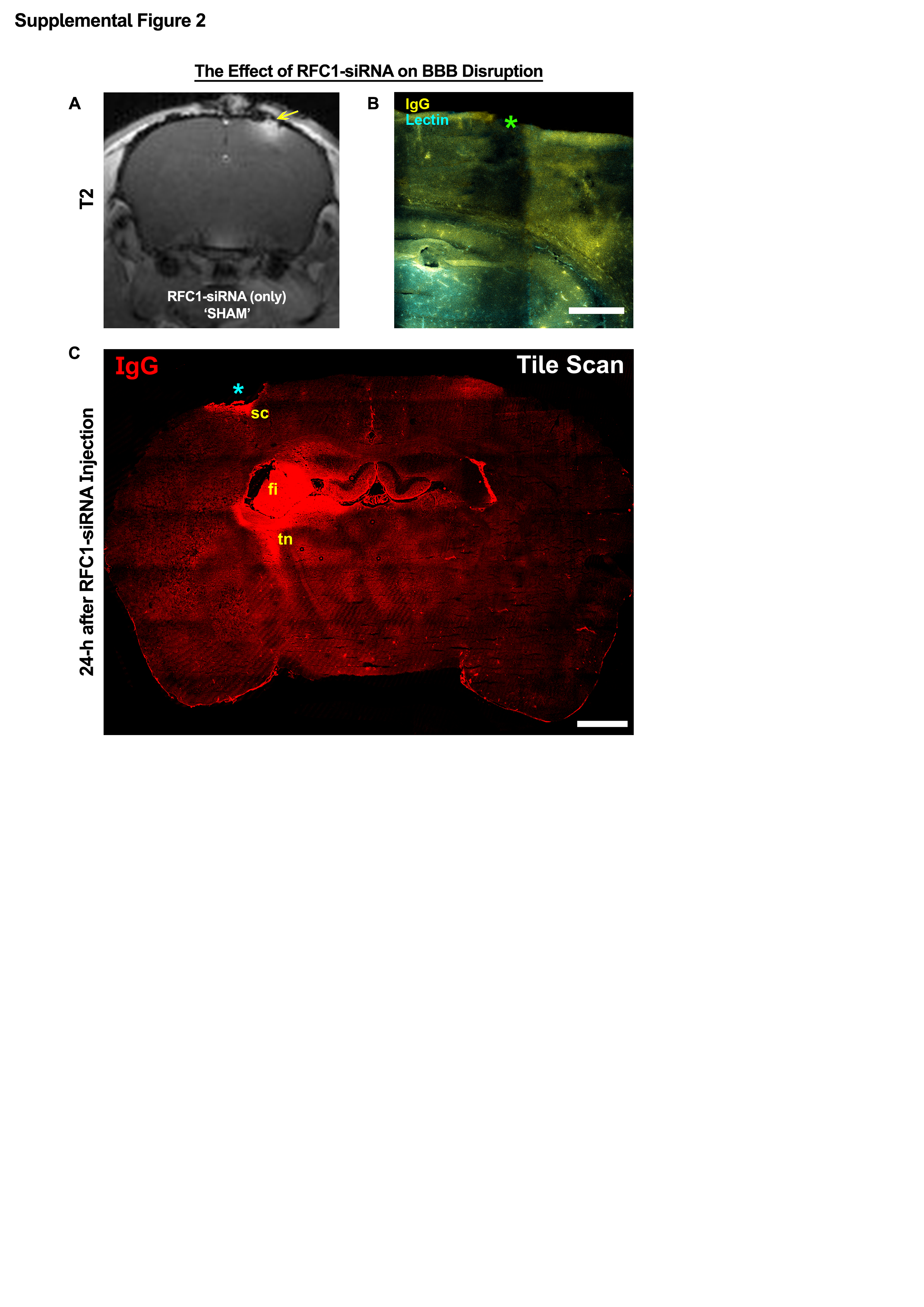

### Supplemental Fig 3_No Layers.tif

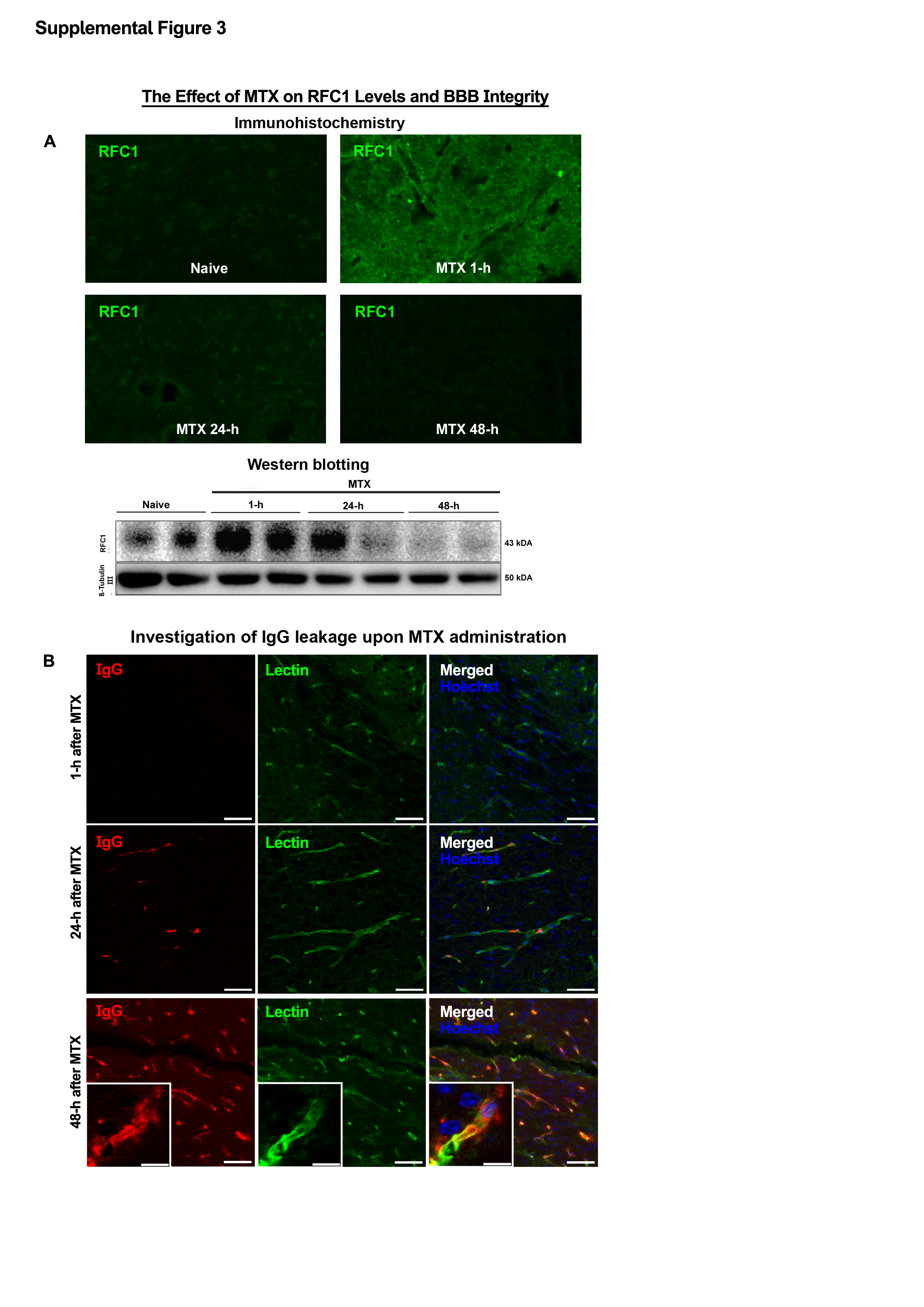

### Supplemental Figure 1.tif

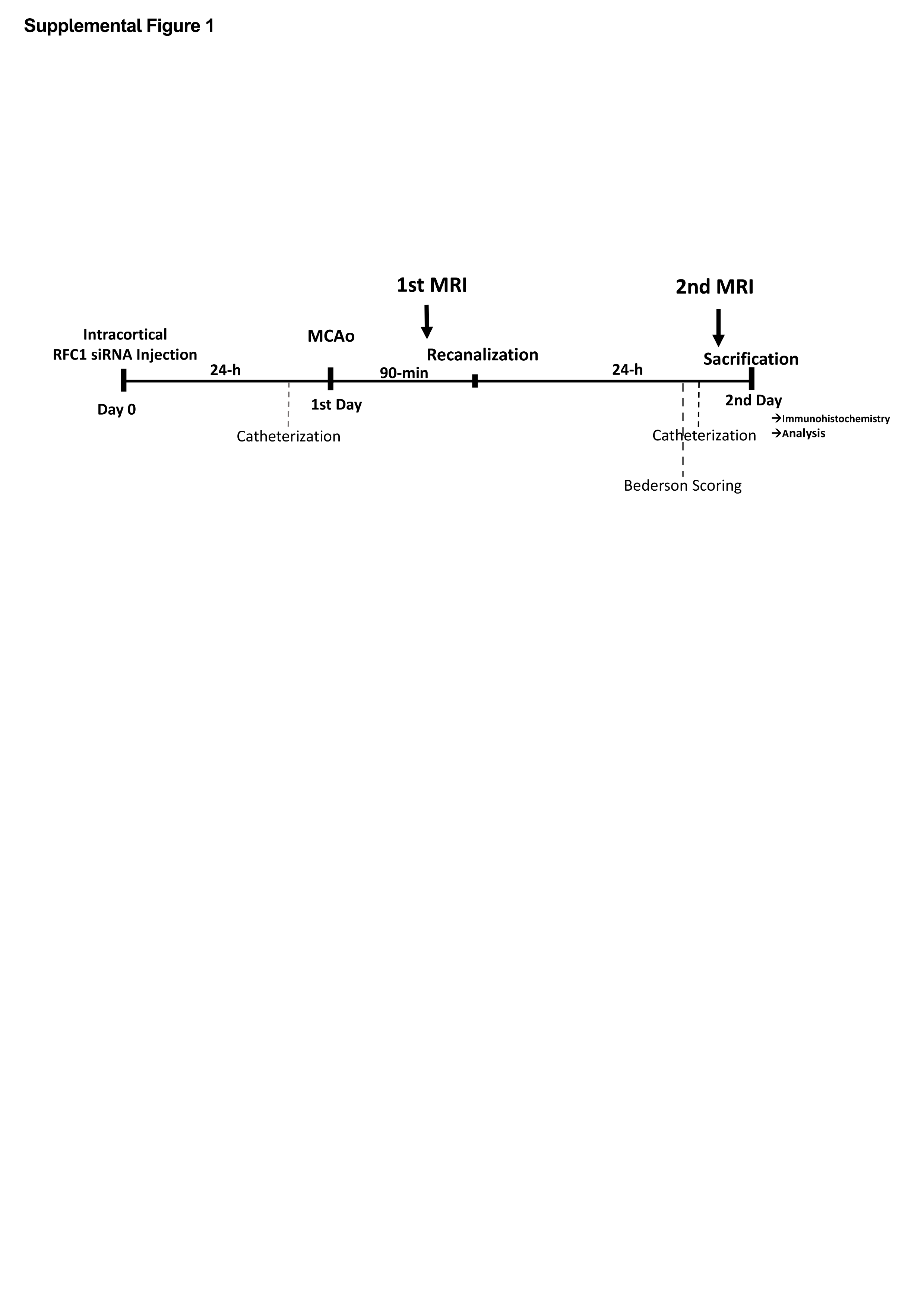
